# Multi-OMIC analysis of brain and serum from chronically-stressed mice reveals network disruptions in purine metabolism, fatty acid beta-oxidation, and antioxidant activity that are reversed by antidepressant treatment

**DOI:** 10.1101/490748

**Authors:** Peter J. Hamilton, Emily Y. Chen, Vladimir Tolstikov, Catherine J. Peña, Punit Shah, Kiki Panagopoulos, Ana N. Strat, Deena M. Walker, Zachary S. Lorsch, Nicholas L. Mervosh, Drew D. Kiraly, Rangaprasad Sarangarajan, Niven R. Narain, Michael A. Kiebish, Eric J. Nestler

## Abstract

Major depressive disorder (MDD) is a complex condition with unclear pathophysiology. Molecular disruptions within the periphery and limbic brain regions contribute to depression symptomatology. Here, we utilized a mouse chronic stress model of MDD and performed metabolomic, lipidomic, and proteomic profiling on serum plus several brain regions (ventral hippocampus, nucleus accumbens, and prefrontal cortex) of susceptible, resilient, and unstressed control mice. Proteomic analysis identified three serum proteins reduced in susceptible animals; lipidomic analysis detected differences in lipid species between resilient and susceptible animals in serum and brain; and metabolomic analysis revealed pathway dysfunctions of purine metabolism, beta oxidation, and antioxidants, which were differentially associated with stress susceptibility vs resilience by brain region. Antidepressant treatment ameliorated MDD-like behaviors and affected key metabolites within outlined networks, most dramatically in the ventral hippocampus. This work presents a resource for chronic stress-induced, tissue-specific changes in proteins, lipids, and metabolites and illuminates how molecular dysfunctions contribute to individual differences in stress sensitivity.

## INTRODUCTION

Major Depressive Disorder (MDD), a complex, heterogeneous syndrome, is the leading cause of disability worldwide. The symptoms of MDD range from emotional and cognitive impairments, including depressed mood, anhedonia, and impaired concentration as well as systemic dysfunctions that affect energy, appetite, cardiovascular function, and immunity. These diverse symptoms suggest the dysregulation of multiple brain regions and peripheral tissues, and there is significant evidence for brain region-specific disruptions in multiple molecular signaling pathways in MDD^1^. Tricyclic antidepressants (TCAs) initially inhibit serotonin and norepinephrine reuptake, but the brain signaling networks affected upon their chronic administration—required for therapeutic efficacy—remain insufficiently understood. Thus, we do not fully grasp the molecular changes that occur throughout the brain and periphery of depressed individuals, and we do not understand the extent to which commonly used antidepressants impinge upon these aberrant molecular mechanisms to improve symptomatology. By more completely understanding the tissue-specific molecular consequences of chronic stress, and illuminating how antidepressants affect these molecular networks, it may be possible to design more targeted MDD treatments.

We utilize an ethologically-validated mouse model of depression called chronic social defeat stress (CSDS), where mice are exposed chronically to a social stress, which induces a range of MDD-like behavioral and molecular changes in a subset (∼50%) of animals, referred to as susceptible^2,3^. These defects are ameliorated by chronic antidepressant treatment^4,5^. The remainder of the stress-exposed population does not display most of these behavioral abnormalities and are referred to as resilient^3^. This divergence in vulnerability to stress is observed within human populations^6^, and inducing these effects within an isogenic mouse population provides the ability to investigate the tissue-specific mechanisms that contribute to individual differences in MDD risk.

In the present study, we employ metabolomic, lipidomic, and proteomic analyses of the ventral hippocampus (vHipp), nucleus accumbens (NAc), and prefrontal cortex (PFC)—all implicated in MDD—and serum samples from susceptible, resilient, and control (undefeated) mice in order to quantify comprehensively the changes that occur within these tissues in response to CSDS. Using this approach, we discover that many of the molecules affected by CSDS are involved in pathways of nucleotide metabolism, fatty acid beta oxidation, and antioxidant function. These pathways are differentially associated with susceptibility or resilience depending on the brain region involved. We also analyzed the effect of chronic administration of imipramine, a standard tricyclic antidepressant which corrects behavioral abnormalities in susceptible mice, on these multi-OMIC endpoints. We observe that many of the same pathways are affected by imipramine treatment, further evidence that activity of these pathways contribute to stress responses. Together, this work provides a rich dataset to explore the tissue-specific, molecular mechanisms that differentiate stress resilient and stress susceptible animals, and outlines strongly-affected protein, lipid, and metabolite pathways that present promising targets for antidepressant drug discovery efforts.

## METHODS AND MATERIALS

### Animals and treatments

Adult male 7-8 week old C57BL/6J mice and 6-month old CD1 retired male breeders (CD1 aggressors) were housed at 22-25°C in a 12-hr light/dark cycle and provided food and water *ad libitum*. All tests were conducted during the light cycle. Experiments were conducted in accordance with the guidelines of the Institutional Animal Care and Use Committee at Mount Sinai.

### Chronic social defeat stress, social interaction testing, and elevated plus maze

We utilized an established CSDS protocol as described previously^2,3^. Briefly, C57BL/6J mice were exposed for ten consecutive days to a novel aggressive CD1 retired breeder for 10 min and were then separated from the aggressor by a perforated divider to maintain constant sensory contact for the remainder of the day. Mice were tested for social interaction (SI) 24 hr after the last social defeat by first allowing 2.5 min for the test mouse to explore an arena containing a plexiglass wire mesh cage centered against one wall of the arena (target absent). In the second 2.5 min test, the same test mouse was returned to the arena with a novel CD1 mouse contained in the plexiglass and wire cage (target present). Based on the social interaction ratio, defined as time spent in the ‘interaction zone’ with target present divided by the time spent with target absent, mice were characterized as susceptible (SI ratio < 1) or resilient (SI ratio > 1). Control mice were housed identically, yet never came in physical or sensory contact with a CD1 aggressor.

For antidepressant experiments, control, resilient, and susceptible populations were treated twice-daily with intraperitoneal (IP) injections of saline or imipramine (10 mg/kg)^5^. Treatment-induced changes in MDD-like behaviors were quantified by re-analyzing social interaction behaviors and performing elevated plus maze analysis. For the latter, mice were tested in a standard maze for 10 min, monitored by Ethovision XT as described previously^7^. Time in the open arms of the plus maze was quantified and expressed as a percent of total time.

### Tissue preparation

vHipp, NAc, and PFC tissue and serum were collected from rapidly euthanized animals, immediately frozen, and stored at -80°C. Tissue samples were pooled from between 7-11 animals depending on brain region, see Figure 1C, and combined in Omni homogenization bead tubes to create ∼20 mg samples. The sera from two animals were pooled to create 400 µL samples. The aggregation of tissue and serum was performed to equalize SI ratios for samples in each group (resilient SI ratio: ∼1.4; susceptible SI ratio: ∼0.8; control SI ratio: ∼1.35). From these pooled samples, all analyses were performed in parallel. Pooled samples were homogenized in water at 4°C in an Omni Bead Ruptor 24 (Omni International, Tulsa, OK) and the protein content of each homogenate were determined via a bicinchoninic assay. Aliquots of 100 µg protein, 10 mg tissue weight, and 0.5 mg protein were separated for proteomics, metabolomics, and structural lipidomics analysis, respectively. Aliquots of pooled serum samples were likewise taken for proteomics, metabolomics, structural lipidomics, and mediator lipidomics analysis.

**FIGURE 1.**
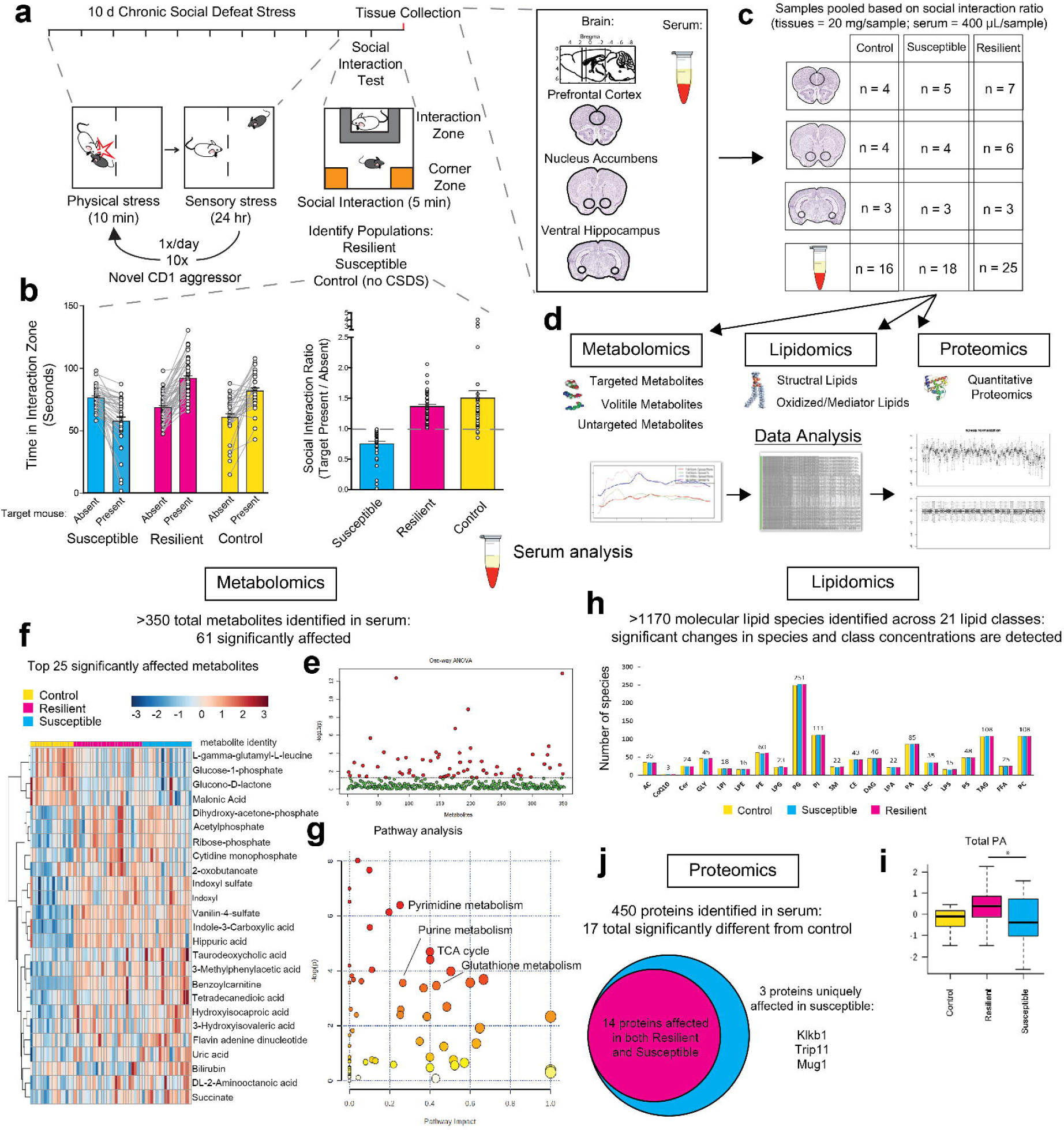
Study overview and metabolomic, lipidomic, and proteomic analysis of serum from resilient and susceptible populations of chronically stressed mice. (**a**) Graphical illustration of workflow for chronic social defeat stress (CSDS) to differentiate mice into susceptible and resilient populations. All tissues harvested for analysis are displayed. (**b**) Social interaction (SI) data from all mice, tested 24 hr after the last CSDS bout. Susceptible animals spend significantly less time in the Interaction Zone (IZ) when the novel CD1 target mouse is present (n = 37; *p* < 0.0001; paired, two-tailed *t* test). Resilient (n = 50) and Control (n = 33) animals spend significantly more time in the IZ when the target mouse is present (*p* < 0.0001; paired, two-tailed *t* test). SI ratio is quantified for each mouse, with resilience as an SI ratio > 1 and susceptibility as an SI ratio < 1. (**c**) To generate sufficient material for parallel analyses, tissues were pooled by SI ratio to achieve 20 mg/sample for brain tissues and 400 μL/sample for serum. (**d**) From this pooled sample, all processing and analysis occurred. (**e**) Metabolomic analysis of serum: 61 important features selected by ANOVA plot with a *p* value threshold of 0.05. (**f**) Heatmap of top 25 affected metabolites shows differences in metabolite levels, localized to experimental groups. (**g**) Pathway analysis of changed metabolites in the serum reveals purine and pyrimidine metabolism, the tricarboxylic acid cycle (TCA cycle), and antioxidant function, among other functions, as significantly affected in the serum of these chronically-stressed mice. (**h**) Lipidomic serum analysis discriminated more than 1170 different lipid species across 21 lipid classes. (**i**) Total circulating levels of phosphatidic acid (PA) were increased in animals resilient to CSDS relative to susceptible animals (*F*_2,58_ = 3.80; *p <* 0.05; Fisher’s LSD). (**j**) Proteomic analysis of serum: a total of 450 proteins were detected, with 17 proteins identified as significantly different from undefeated control animals. Of these 17, three proteins were significantly decreased solely in the susceptible cohort: kallikrein B1 (Klkb1), murinoglobulin-1 (Mug1), and thyroid receptor-interacting protein 11 (Trip11).

### Targeted and untargeted metabolomic analysis

Metabolomic analyses were performed using untargeted and targeted protocols as described previously^8-11^. Metabolite extraction was achieved using a mixture of isopropanol:acetonitrile:water (3:3:2 v/v/v). Tissue samples were homogenized in an extraction mixture using Fisherbrand™ Model 120 Sonic Dismembrator. Extract analysis was performed using GC/MS, RP-LC/MS, and HILIC-LC/MSMS protocols as described^8^. Quality control was performed using metabolite standards mixture and pooled samples. Collected raw data were manually inspected, merged, and imputed. Statistical analysis was performed with MetaboAnalyst 3.0^12^. Metabolite Set Enrichment Analysis (MSEA) was used to interrogate functional relations, which describes the correlation between compound concentration profiles and clinical outcomes.

### Structural lipidomic analysis

A cocktail of deuterium-labeled and odd chain phospholipid standards from diverse lipid classes was added to 25 µL of thawed serum or tissue homogenate with 0.5 mg protein as measured via a bicinchoninic assay. Standards were chosen so that they represented each lipid class and were at designated concentrations chosen to provide the most accurate quantitation and dynamic range for each lipid species. 4 mL chloroform:methanol (1:1, v/v) was added to each sample and the lipid extraction was performed as described^13,14^. Lipid extraction was automated using a customized sequence on a Hamilton Robotics STARlet system (Hamilton, Reno, NV) to meet the high-throughput requirements. Lipid extracts were dried under nitrogen and reconstituted in 68 µL chloroform:methanol (1:1, v/v). Samples were flushed with nitrogen and stored at -20°C. Samples were diluted 50 times in isopropanol:methanol:acetonitrile:water (3:3:3:1, by volume) with 2 mM ammonium acetate in order to optimize ionization efficiency in positive and negative modes. Electrospray ionization-MS was performed on a TripleTOF® 5600^+^ (SCIEX, Framingham, MA), coupled to a customized direct injection loop on an Ekspert microLC200 system (SCIEX) as described^7^. Lipid species with >50% missing values were removed, with remaining missing values estimated (the half of the minimum positive values in the original data assumed to be the detection limit) and IQR filtered, normalized to the median, glog transformed, and autoscaled. One-way ANOVA and volcano plot analysis was performed using Metaboanalyst 4.0^15^.

### Mediator lipidomic anaylsis

A mixture of deuterium-labeled internal standards was added to aliquots of 100 µL serum, followed by 3x volume of sample of cold methanol (MeOH). Samples were vortexed for 5 min and stored at -20°C overnight. Cold samples were centrifuged at 14,000g at 4°C for 10 min, and the supernatant was then transferred to a new tube and 3 mL of acidified H_2_O (pH 3.5) was added to each sample prior to C18 SPE columns (Thermo Pierce) and performed as described^16^. The methyl formate fractions were collected, dried under nitrogen, and reconstituted in 50 µL MeOH:H_2_O (1:1, v/v). Samples were transferred to 0.5 mL tubes and centrifuged at 20,000g at 4°C for 10 min. Thirty-five µL of supernatant were transferred to LC– MS vials for analysis using the BERG LC–MS/MS mediator lipidomics platform as described^17^.

### Proteomic Analysis

Sixty-five µL of serum were delipidated using Lipisorb and then depleted using a multiple affinity removal spin cartridges Mouse-3 (Agilent Technologies). Low abundant proteins were collected in 100% Agilent Buffer A. Delipidated and depleted samples were used for determination of protein concentration using a Coomassie Bradford Protein Assay Kit (Thermo Pierce). Tissues were lysed using 7M urea, 2M thiourea, 1% Halt Protease and Phosphatase Inhibitor cocktail and 0.1% SDS, followed by sonication. After lysis, samples were centrifuged, and supernatant was used for proteomics analysis. The protein concentration was determined using Coomassie Bradford Protein Assay Kit. Proteins were reduced in 10 mM Tris(2-carboxyethyl) Phosphine (TCEP) for 30 min at 55°C and alkylated in 18.75 mM iodoacetamide for 30 min at room temperature in the dark. Proteins were precipitated overnight using acetone. Protein pellets were reconstituted in 200 mM tetraethylammonium bicarbonate (TEAB) and digested with trypsin at 1:40 (trypsin:protein) overnight at 37°C. Peptides were then labeled with Tandem Mass Tag (TMT) 10-plex isobaric label reagent set (Thermo Pierce) using manufacturer’s protocol. Labeling reaction was quenched with 5% hydroxylamine for 15 min before being combined into each respective multi-plex (MP). Pooled samples were dried in a vacuum centrifuge followed by desalting using C-18 spin columns (Thermo Pierce). The eluate from C-18 was dried in a vacuum centrifuge and stored at -20°C until LC-MS/MS analysis.

LC-MS/MS analysis was performed using a Waters nanoAcquity 2D LC system coupled to a Thermo Q Exactive Plus MS. TMT-labeled samples were fractionated online into 12 basic reverse phase fractions. Each fraction was subjected to 90 min reverse phase separation. Data-dependent Top-15 acquisition method was used for MS analysis. Parameters used for Q-Exactive plus were full MS survey scans at 35,000 resolution, scan range of 400-1800 Thompsons (Th; Th = Da/z). MS/MS scans were collected at a resolution of 35,000 with a 1.2 Th isolation window. Only peptides with charge +2, +3, and +4 were fragmented with a dynamic exclusion of 30 sec.

Raw LC-MS/MS data were then processed using Proteome Discoverer v1.4 (Thermo) by searching a Swissport Mouse database using the following parameters for both MASCOT and Sequest search algorithms: tryptic peptides with at least six amino acids in length and up to two missed cleavage sites, precursor mass tolerance of 10 ppm, fragment mass tolerance of 0.02 Da; static modifications: cysteine carbamidomethylation, N-terminal TMT10-plex; and dynamic modifications: asparagine and glutamine deamindation, methionine oxidation, and lysine TMT10-plex.

## RESULTS

### Widespread molecular changes are induced by chronic social defeat stress

To resolve the diverse molecular changes that occur within the brain and serum of mice in response to chronic stress, we exposed C57BL/6J mice to ten days of CSDS, and 24 hr after the final defeat we performed a social interaction (SI) test to identify populations of mice that are either susceptible or resilient to the stress (**Fig. 1a,b**). We then performed metabolomic, lipidomic, and proteomic profiling in parallel on pooled samples of serum and of vHipp, NAc, and PFC of susceptible, resilient, and undefeated control mice to capture the full spectrum of molecular changes in blood and brain that differentiate these populations (**Fig. 1c,d**).

In serum, metabolomic analysis identified more than 350 metabolites, with 61 significantly affected as assessed by one-way ANOVA with a p value threshold of -log_10_(p)>1.3 (**Fig. 1e**). The identities of these metabolites along with post-hoc analyses are presented in **Supplemental Table 1a**. The top 25 supervised heatmap shows localized group differences in metabolite levels (**Fig. 1f**). Pathway analysis identified many of the affected metabolites as important in the biological functions of nucleotide metabolism, as both purine and pyrimidine pathways were identified, as well as energy production and antioxidant function, in the form of glutathione metabolism (**Fig. 1g**). Lipidomic analysis identified greater than 1170 unique lipid species across 21 lipid classes, with no significant group differences in the number of identifiable lipid species (**Fig. 1h**). However, we detected significant changes in oxidized lipid mediators, as well as changes in the concentration of 57 structural lipid species; for example, total serum levels of phosphatidic acid (PA) were significantly increased in resilient animals (**Fig. 1i,** list of structural lipid species in **Supp. Table 1b;** list of 9 oxidized/mediator lipids affected in **Supp. Table 1c**). Lastly, proteomic analysis detected 450 unique proteins in serum, with a total of 17 identified as significantly different from control animals (**Supp. Table 1d**). Three proteins were significantly decreased exclusively in the serum of susceptible animals: kallikrein B1 (Klkb1), murinoglobulin-1 (Mug1), and thyroid receptor-interacting protein 11 (Trip11) (**Fig. 1j**). Collectively, these comprehensive analyses differentiate the molecular composition of the periphery of susceptible vs resilient animals.

Stress-responses are mediated through the functions of several brain regions^18^. To investigate how chronic stress impacts the molecular composition of key brain regions, we performed multi-OMIC analyses on vHipp, NAc, and PFC tissues. With metabolomic analysis in the vHipp, an unsupervised heatmap of all detected metabolites clustered according to group, indicating substantial group differences in general metabolite composition (**Fig. 2a**). With the exception of one sample, control and susceptible groups were the most metabolically different, with the resilient group occupying an intermediate metabolic identity. We identified 39 significantly altered metabolites in this brain region (**Fig. 2b,** identities in **Supp. Table 2a**). A supervised heatmap of the top 25 affected metabolites shows dramatic concentration differences between control, susceptible, and resilient groups, with many of the affected metabolites being purine nucleotides, carnitine donors and modified fatty acids involved in beta oxidation, and antioxidants (**Fig. 2c**). The same metabolomic detection analyses performed in the NAc revealed 19 significantly altered metabolites (**Fig. 2d,** identities in **Supp. Fig. 2b**), whereas in the PFC only seven significant metabolites were detected (**Fig. 2e,** identities in **Supp. Fig. 2c**), revealing a different metabolic impact of chronic stress by brain region.

**FIGURE 2.**
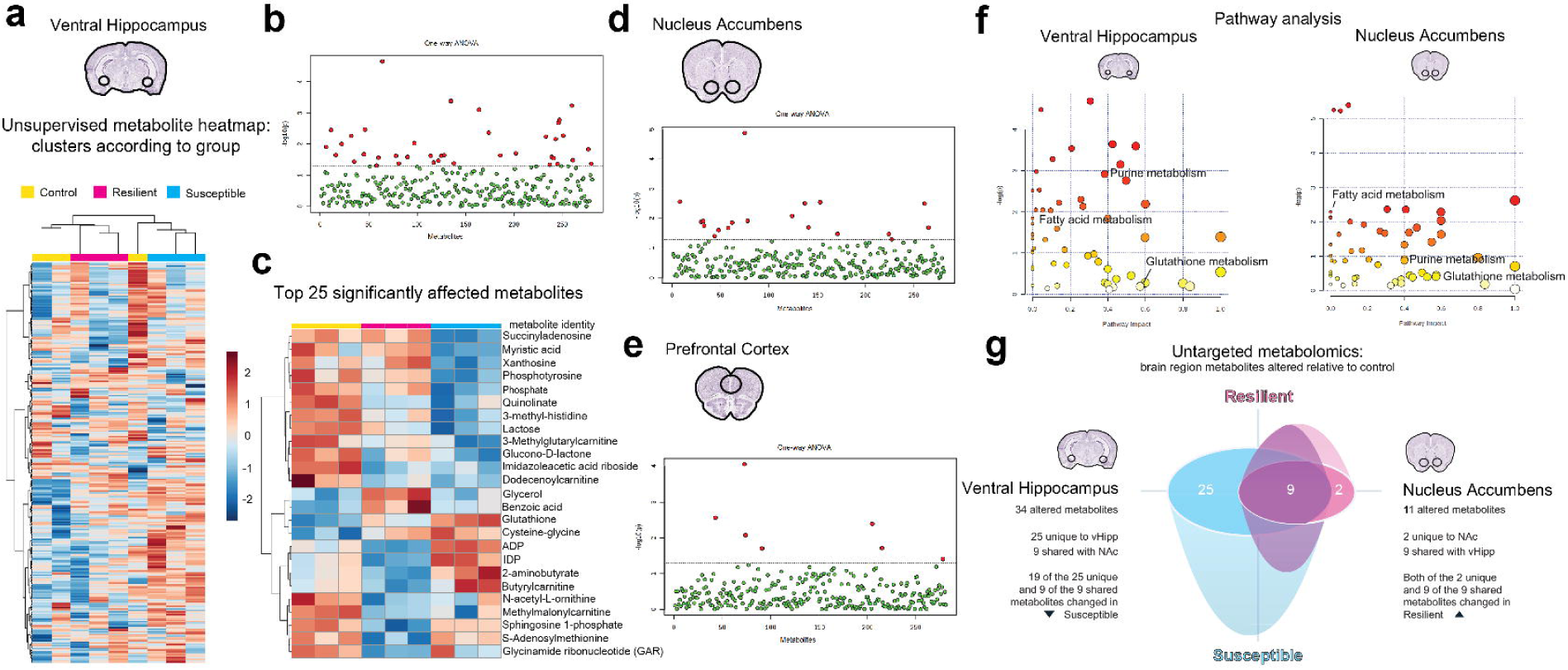
Metabolites involved in purine metabolism, beta oxidation of fatty acids, and antioxidant function are differentially affected by chronic stress in each brain region. (**a**) Ventral hippocampus (vHipp) samples show robust metabolic changes after CSDS. Unsupervised clustering of metabolite heatmaps largely aggregate by group, revealing substantial differences in metabolic composition that differentiate control, resilient, and susceptible animals. (**b**) 39 significantly changing metabolites identified in the vHipp by ANOVA plot with a *p* value threshold of 0.05. (**c**) A supervised heatmap of the top 25 significantly affected metabolites reveal dramatic differences in metabolite concentrations in each group. (**d**) 19 significantly changing metabolites identified in the nucleus accumbens (NAc) by ANOVA plot with a *p* value threshold of 0.05. (**e**) 7 significantly changing metabolites identified in the prefrontal cortex (PFC) by ANOVA plot with a *p* value threshold of 0.05. (**f**) Pathway analysis of changed metabolites in the vHipp and NAc reveals purine metabolism, beta oxidation, and antioxidant function as significantly affected in both of these brain regions. (**g**) Untargeted metabolomic analysis independently validates that the vHipp is the most stress-affected brain region with 34 significantly altered metabolites. In the NAc, 11 affected metabolites are detected, nine of which are affected in both the vHipp and the NAc. However, metabolic changes in the NAc are associated with resilience to CSDS, while metabolic changes in the vHipp are associated with susceptibility to CSDS, since all metabolites in the NAc are solely altered in resilient animals, whereas a majority of the metabolites in the vHipp are altered in susceptible animals.

Several of the same or similar metabolites were affected in all brain areas. To investigate this further, we performed pathway analysis in the two most dynamic brain regions, the vHipp and NAc (**Fig. 2f**). In both regions, many of the same molecular pathways were affected, including purine metabolism, fatty acid metabolism, and metabolism of the antioxidant glutathione, revealing a common impact of CSDS on these pathways across brain regions.

To understand whether the effect of CSDS on these various pathways was associated with susceptibility or resilience by brain region, and to independently validate the identified metabolites from our earlier analysis, we performed untargeted metabolomic profiling in parallel to our targeted analyses. This separate analysis validated our previous metabolite profiling in that many of the same or similar molecules were identified *de novo* (**Supp. Table 3**). Relative to undefeated controls, untargeted metabolomics identified 34 metabolites significantly altered in the vHipp, and 11 metabolites significantly altered in the NAc (**Fig. 2g**), again revealing a different metabolic response to stress by brain region. While the NAc and vHipp overlapped in a considerable number of affected metabolites, in vHipp the regulation of identified molecules occurred primarily in susceptible animals (28 of 34 metabolites altered in susceptible mice only), whereas in the NAc alterations occurred entirely in resilient animals (11 of 11 metabolites altered in resilient only). Thus, while these brain regions utilize many of the same molecular pathways in response to chronic stress, metabolic impact in the NAc is associated with resilience, with metabolic and impact in the vHipp associated with susceptibility to chronic stress. With lipidomic analysis, we observed 36 altered lipid species in the vHipp with multiple species of long chain phosphatidylethanolamine (PE), a major constituent of the myelin sheath, being decreased in susceptible animals, as well as notable regulation of sphingolipids and cardiolipin species (**Supp. Table 4a**). We identified 54 altered lipid species in the NAc (**Supp. Table 4b**), with 28 lipid species altered in the PFC (**Supp. Table 4c**). With proteomic analysis, we observed 51 altered proteins in the vHipp (**Supp. Table 5a**), 53 in the NAc (**Supp. Table 5b**), and 71 in the PFC (**Supp. Table 5c**). These data collectively provide a rich resource to explore the brain-region specific, molecular adaptations induced by chronic stress.

### Imipramine treatment regulates key molecules in the brain and periphery

To investigate the molecular effects of imipramine treatment, we again stratified C57BL/6J mice into resilient and susceptible populations after CSDS and divided animals into their future treatment groups by equalizing SI ratios (**Fig. 3a,b**). Susceptible, resilient, and control mice were treated twice daily for 14 days with an IP injection of either saline or imipramine (10 mg/kg) (**Fig. 3a,c**). Twenty four hr after the last injection, all mice were examined in a second social interaction test, as well as an elevated plus maze test, to determine the effect of imipramine exposure.

**FIGURE 3.**
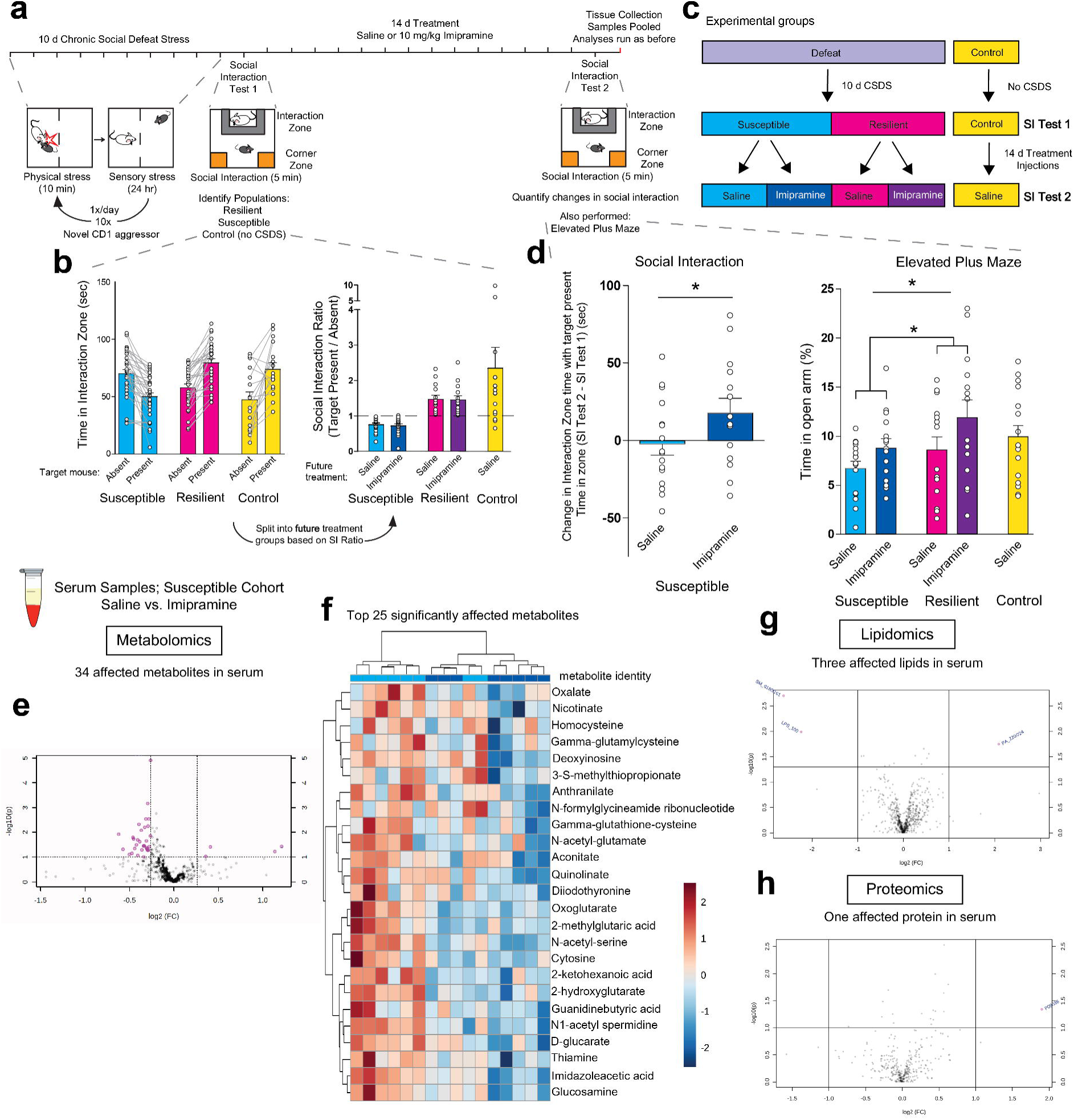
Chronic imipramine treatment improves susceptible behaviors and correlated peripheral molecular profiles. (**a**) Graphical illustration of workflow for CSDS, treatment schedule, and behavioral testing. Serum, vHipp, NAc, and PFC were again harvested, pooled by SI ratio, and analyzed as before (see Fig. 1a). (**b**) SI test 1 data from all mice used to determine susceptible, resilient, and control populations. Susceptible animals spend significantly less time in the IZ when the novel CD1 target mouse is present (n = 33; *p* < 0.0001; paired, two-tailed *t* test). Resilient (n = 28) and Control (n = 16) animals spend significantly more time in the IZ when the target mouse is present (*p* < 0.0001; paired, two-tailed *t* test). SI ratio is quantified for each mouse, and mice are divided into future treatment groups based on equalized average SI ratios per population. (**c**) Representation of workflow to generate each treatment group. (**d**) Post-treatment behavioral data including SI test 2 and elevated plus maze (EPM). Left: SI test 2 in susceptible population quantifying change in time spent in the interaction zone when the novel CD1 target mouse was present between SI test 1 and SI test 2 (n = 14-16; *=*p* < 0.05 by Student’s *t* test). Right: EPM test quantifying anxiety-like behaviors in all groups. Resilient animals spend more time exploring the open arms than susceptible animals (*F*_1,56_ = 4.19; *=*p* < 0.05 by two-way ANOVA), and treatment significantly affects anxiety-like behaviors (*F*_2,56_ = 4.93; *=*p* < 0.05 by two-way ANOVA). (**e**) Metabolomic analysis of serum from imipramine-treated susceptible cohort. Volcano-plot highlights 34 important features selected by ANOVA plot. (**f**) Unsupervised heatmap of top 25 affected metabolites largely clusters according to treatment and demonstrates prevailing imipramine-mediated down-regulation of key metabolites, many of which are involved in energy homeostasis and antioxidant function. (**g**) Volcano plot of the three significantly affected lipids. PA = phosphatidic acid, SM = sphingomyelin, LPS = lysophosphatidylserine. (**h**) Volcano plot reveals one affected protein. P09036 = Spink1.

As expected^5^, chronic imipramine administration reversed social interaction deficits seen in susceptible mice. We quantified the difference in time that the animal spent in the interaction zone when the target CD1 mouse was present between SI test 1 (before treatment) and SI test 2 (after treatment). While saline treatment had a minimal effect on the change in interaction time, there was a significant improvement in imipramine-treated susceptible mice (**Fig. 3d**).

We next performed the elevated plus maze assay. We observed a significant difference in time spent on the open arm between susceptible and resilient groups, with the former group spending less time in the open arm, indicating that group phenotypes persisted throughout the treatment window. We also observed a significant treatment effect in both susceptible and resilient groups with imipramine treatment increasing the amount of time spent exploring the open arm (**Fig. 3d**). Collectively, these data indicate that imipramine improves behavioral abnormalities induced by CSDS. To understand the molecular pathways affected by imipramine exposure in chronically-stressed mice, we performed multi-OMIC profiling on pooled samples of serum and of vHipp, NAc, and PFC of all mice.

In the serum of susceptible animals treated with saline or imipramine, metabolomic analysis identified 34 affected metabolites with a p value threshold of -log_10_(p)>1 (**Fig. 3e,** identities in **Supp. Table 6**). An unsupervised heatmap representing the top 25 significantly affected metabolites shows partial clustering in serum metabolite composition by treatment (**Fig. 3f**). Notably, imipramine treatment reduces the levels of many metabolites, with many of the affected metabolites being involved in previously identified functions, including energy homeostasis and antioxidant function. This indicates the ability of imipramine to reverse certain CSDS-induced metabolic derangements in peripheral blood in concert with its antidepressant effect.

With lipidomic analysis, we observed that imipramine treatment significantly affected the concentrations of three lipids: sphingomyelin and lysophopshatylserine species were decreased (**Fig. 3g**), and there was an increase in a PA species, which is consistent with the earlier-observed increase in PA species specifically in resilient animals.

Lastly, with proteomic analysis, we identified one significantly affected protein, Spink1 (**Fig. 3h**). Spink1 acts as a serine-protease inhibitor, which would have opposite regulatory actions relative to the earlier-identified susceptible-specific protein, Klkb1 (**Fig. 1j**), which is itself a serine-protease. This observation highlights the potentially important role for peripheral protease activity in contributing to stress responses, and is supported by the observation that elevated levels of the serine protease prostasin were observed in urine of patients with MDD^19^ and that alpha-1-antitrypsin, a protease inhibitor, is differentially affected in the serum of individuals with a current MDD diagnosis relative to remitted MDD^20^.

We next performed metabolomic analysis in each brain region. In the vHipp, as in serum, we observed primarily a down-regulation of key metabolites, with 18 total metabolites affected (**Fig. 4a,** list in **Supp. Table 7a**). A heatmap of the top 25 affected metabolites clusters by treatment, revealing how, of all brain regions, imipramine treatment most concisely affects the vHipp and acts to diminish the levels of metabolites that were previously observed to be elevated in the vHipp of susceptible animals, including sphingosine 1-phospate and metabolites involved in antioxidant function and *de novo* purine synthesis (**Fig. 4a**). In the NAc, 11 metabolites are affected, with a shift towards up-regulation of metabolite levels by imipramine treatment, and the heatmap of the top 25 affected metabolites less clearly separates by treatment (**Fig. 4b,** list in **Supp. Table 7b**). Lastly, only three metabolites were affected in the PFC (**Fig. 4c,** list in **Supp. Table 7c**). These data mirror the observations of **Figure 2**, in that the brain areas that were most affected metabolically by stress are also most affected by imipramine treatment. Taken together, these data show that imipramine reversal of CSDS-induced behavioral deficits is most associated with metabolic effects in the vHipp.

**FIGURE 4.**
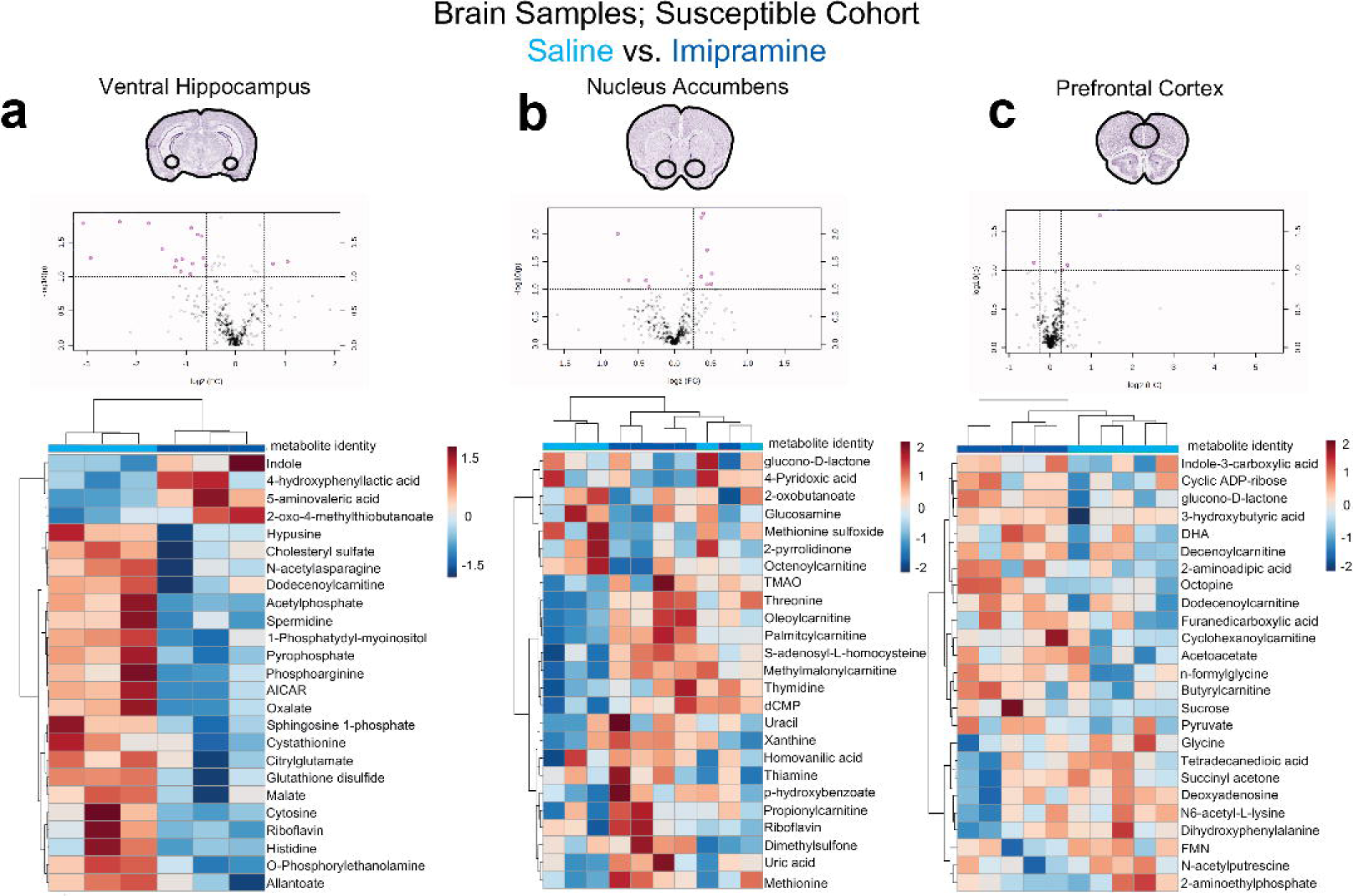
Imipramine-induced molecular adaptations by brain region within susceptible animals. (**a**) Top: Volcano plot of 18 changing metabolites identified in the vHipp. Bottom: Heatmap of top 25 affected metabolites clusters by treatment, revealing discrete metabolic profiles induced by imipramine. Saline treatment is light blue, imipramine treatment is dark blue (**b**) Top: Volcano plot of 11 changing metabolites identified in the NAc. Bottom: Heatmap of top 25 affected metabolites. (**c**) Top: Volcano plot of three changing metabolites identified in the PFC. Bottom: Heatmap of top 25 affected metabolites.

### Metabolic network construction

We utilized the metabolomics data in serum and brain to construct the net effect of CSDS and imipramine treatment on metabolic networks. In the vHipp, we observed consistent changes in many of the metabolites within the *de novo* purine synthesis pathway with resilient animals showing decreases and susceptible animals showing increases in these metabolites (**Fig. 5a**). Furthermore, many of the metabolites of purine breakdown are decreased in susceptible animals, whereas ADP, a precursor of ATP, is significantly elevated in susceptible and decreased in resilient animals. Importantly, a key product of ATP, s-adenosyl methionine (SAM-e) is diminished in the vHipp of resilient animals. SAM-e is a major methyl donor used for protein and lipid methyation, and differences in its bioavailability within the vHipp could have sweeping consequences on cellular functions. Further, SAM-e is a substrate for the transsulfuration pathway, producing glutathione and other antioxidant molecules, which we consistently observed to be elevated in susceptible animals and down-regulated in serum and vHipp of these susceptible animals upon imipramine treatment.

**FIGURE 5.**
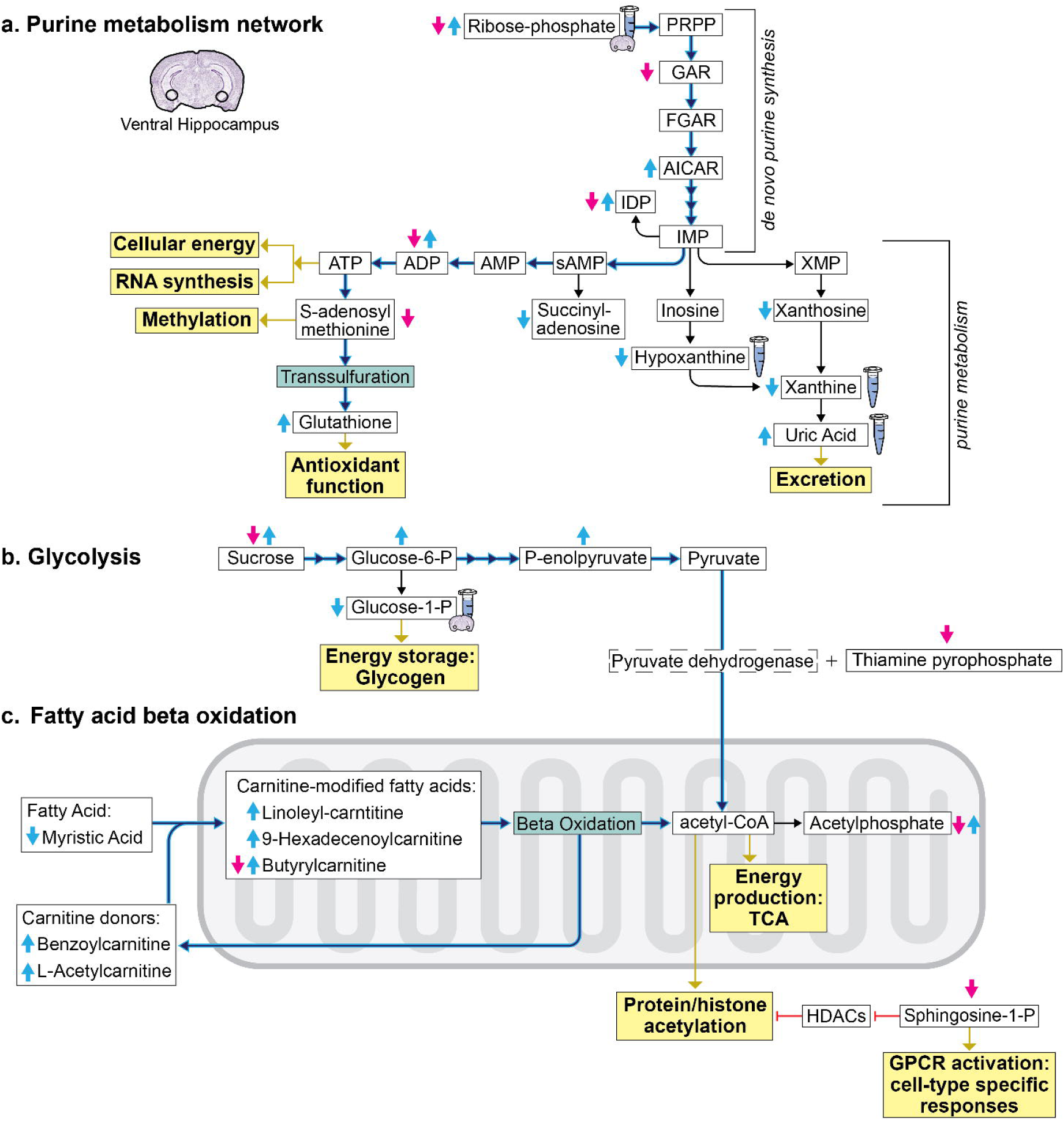
Ventral hippocampus metabolic networks differentially affected in susceptibility vs resilience. Red and blue arrows indicate regulation of metabolite levels observed from our datasets in resilient and susceptible animals, respectively. The hyperactive pathway through these networks experienced by susceptible animals is highlighted in blue. Metabolite relationships were derived from KEGG database and the IUBMB-Sigma-Nicholson Metabolic Pathways Chart. Solid black arrows represent direct metabolic reactions. Yellow boxes represent basic cellular functions affected by key metabolites. PRPP: phosphoribosyl pyrophosphate; GAR: glycinamide ribonucleotide; FGAR: 5′-Phosphoribosyl-N-formylglycineamide; AICAR: 5-Aminoimidazole-4-carboxamide ribonucleotide; IMP: inosine monophosphate; IDP: inosine diphosphate; XMP: xanthosine monophosphate; sAMP: succinyl adenosine monophosphate; AMP: adenosine monophosphate; ADP: adenosine diphosphate; ATP: adenosine triphosphate

In the vHipp, beyond the purine network, susceptible and resilient animals show opposing effects in glycolysis and fatty acid beta oxidation pathways. Both pathways are major sources of cellular energy through production of acetyl-CoA for the tricarboxylic acid cycle. Susceptible animals have elevated levels of precursor molecules necessary for the synthesis of pyruvate (**Fig. 5b**). The cofactor thiamine pyrophosphate (TPP), which potentiates the conversion of pyruvate to acetyl-CoA, trends towards decreased in resilient animals. Similarly, susceptible animals show hyper-function of the fatty acid beta oxidation cycle. Beta oxidation is the process through which fatty acids are internalized in the mitochondria via carnitine modification and consumed to yield acetyl-CoA^21^. These susceptible-specific effects in glycolysis and beta oxidation converge in the accumulation of acetyl-CoA. This provides substantially more reactants for the tricarboxylic acid cycle and ATP synthesis, further implicating hyper-energetics in the vHipp of susceptible animals. Also, acetyl-CoA is a source for protein acetylation, including histones^22^. In direct contrast to the susceptible animals, protein acetylation may be decreased in the vHipp of resilient animals via decreases in sphingosine-1-phosphate (S1P) levels. S1P performs many cellular functions, including acting as a direct inhibitor of class I and II histone deacetylases (HDACs)^23^. This potential regulation of histone acetylation levels is consistent with a literature in both mice and rats describing changes in hippocampal histone acetylation in response to stress and antidepressant exposure^5,24-26^.

We were unable to resolve significant differences in ATP or acetyl-CoA themselves, likely due to the fact that these metabolites are consumed rapidly and utilized for numerous cellular functions. However, resolving the full metabolic network, including precursor and product molecules, serves to clearly demonstrate metabolic demands. Taken together, these network analyses demonstrate that the brain-region specific cellular functions of energy production and the global post-translational modifications of methylation and acetylation are likely hyperactive in the vHipp of susceptible animals. This illuminates that individual differences in these cellular functions by brain-region may underlie the divergence in stress response.

## DISCUSSION

Here, we generated a comprehensive dataset that quantifies the protein, lipid, and metabolite changes that occur across the brain and serum of chronically-stressed and imipramine-treated mice. This is a uniquely-detailed dataset of this nature, from which many fundamental discoveries relating to the biological mechanisms underlying stress responses and antidepressant action can be derived. Analyzing these data, we discovered that chronic stress selectively affects the molecular pathways of purine metabolism, beta oxidation, and antioxidant function and that such changes associate with individual differences in behavioral responses to stress, depending on the brain region in which the adaptations occur.

We observed that the three brain regions studied display marked differences in metabolic responses to chronic stress: the vHipp is the most robustly affected, followed closely by the NAc, with the PFC not as affected by chronic stress by these metrics. By contrast, the PFC of mice and humans is affected dramatically by chronic stress and MDD, respectively, at the transcriptional level^1,27^, raising the interesting possibility that different brain regions may undergo different modes of biological regulation under these conditions. Considering that the vHipp is instrumental in processing both depression- and anxiety-like behavior^28,29^, it is not surprising that this region is so heavily affected by chronic stress at the metabolic level. Our findings further illuminate that a major difference between stress-resilient and -susceptible mice is the activity of specific molecular pathways within this brain region (**Fig. 5**). Susceptibility may manifest from metabolic processing to support the hyperactivity of key cellular functions, such as ATP synthesis and availability of molecular reactants for protein post-translational modifications. In support of this notion, treating stressed animals with imipramine down-regulated these molecular substrates in the vHipp and improved CSDS-induced behavioral abnormalities. Resilience is associated with the absence of molecular reactants within the vHipp. In contrast to vHipp, hyperactivity of largely the same molecular pathways within the NAc, a brain region instrumental in processing reward, is correlated with resilience. Therefore, we can conclude that the outlined molecular pathways are selectively sensitive to chronic stress, but that the brain region in which altered function occurs differentiates the animal’s stress susceptibility or resilience.

Metabolites involved in glycolysis, beta oxidation, and the tricarboxylic acid cycle were also consistently affected in serum and brain. Interestingly, imipramine treatment down-regulated key energy-related molecules in the vHipp and up-regulated others in the NAc. For example, diiodothyronine, a thyroid hormone that increases ATP synthesis by blocking feedback inhibition of cytochrome c oxidase^30^, was decreased in vHipp of susceptible animals treated with imipramine (**Supp. Table 7a**). Conversely, many carnitines central to the function of beta oxidation were upregulated in the NAc of susceptible animals treated with imipramine (**Supp. Table 7b**). The availability of cellular energy would affect the functioning of these brain regions. Indeed, reduced neuronal activity in the vHipp is observed in animals resilient to CSDS, with its optogenetic activation promoting susceptibility^31^. Moreover, these observations suggest that imipramine might induce its therapeutic effect, in part, by regulating brain-region specific energy production, and highlights amenable molecular pathways to target with novel antidepressants.

A further proxy of the signaling properties of these brain regions are the region-specific profiles of neurotransmitters. Within the vHipp, resilient animals have reductions in dihydroxyphenylalanine (L-DOPA; **Supp. Table 2a**), a precursor of the neurotransmitter dopamine, suggesting reduced dopaminergic input activity in the vHipp of resilient animals, which would be expected to alter hippocampus function. Also, γ-aminobutyric acid (GABA) is decreased in the vHipp of susceptible animals, possibly indicating a loss in local inhibitory GABAergic tone. As well, there is a trend for decreased 5-hydroxyindole acetic acid, the terminal product of serotonin (5-HT) catabolism, potentially reflecting lowered 5-HT levels in the vHipp of susceptible animals. Serotonergic afferents release 5-HT in the vHipp upon stress exposure and mediate a counter-adaptive mechanism to stress response^32^. 5-HT inhibits the consolidation of stress-related memories^33^, and rodents bred for high anxiety-like behavior have lower levels of stress-induced hippocampal serotonin release^34^. Thus, lowered 5-HT tone within the vHipp would likely contribute to the phenotype of susceptible animals. These data reveal the impact of stress on neurotransmitter synthesis in the vHipp, and are consistent with our findings across multiple levels of molecular analysis that hyperfunction of the vHipp is associated with susceptibility and hypofunction is associated with resilience.

Lipids present another promising class of molecular targets for novel antidepressants. Lipids play key roles in cell membrane function, neurotransmitter synthesis, and cell signaling, and their dysfunction is increasingly appreciated as contributing to MDD and related disorders^35^. We observe carnitines involved in fatty acid beta oxidation as the most dramatically and consistently affected class of molecule across brain regions. In the vHipp, long chain species of PE were depleted in susceptible animals, a potential indication of demyelination as well as of reactive oxygen species. There is evidence that PE methylation is the primary consumer of SAM-e, and in instances of PE depletion, histone proteins replace PE as the primary methyl sinks^36^. Considering that resilient animals have reductions in SAM-e levels in vHipp, and susceptible animals have depletions in PE lipids, this evidence converges to suggest that methylation, and specifically histone methylation, might be oppositely regulated in this brain region of susceptible vs resilient mice.

S1P was also diminished in the vHipp of resilient animals, with imipramine treatment reducing S1P levels in susceptible animals. S1P signals through G-protein coupled receptors^37^, and has been demonstrated to directly inhibit class I and II HDACs^23^. S1P has been reported to promote stress-induced anxiety-like behavior in rats when locally infused into the cerebral ventricles^38^. Thus, lowering levels of S1P in susceptible mice may lessen cell-signaling responses and reduce bulk histone acetylation levels, which would have broad transcriptional consequences and might contribute to imipramine’s therapeutic effects. Previous studies examining the beneficial actions of imipramine have linked its antidepressant effects to reductions in ceramide levels^35^. However, this is the first demonstration that a further metabolite of ceramide, namely, S1P, is associated with imipramine’s antidepressant actions.

In serum, we detected many stress-induced alterations in levels of lipid species. In resilient animals, several PA species were elevated, and the only up-regulated lipid in response to antidepressant treatment was a long-chain PA (**Fig. 3g**). PA is involved in phospholipid biosynthesis, membrane remodeling, and second messenger signaling, and regulates immune response by stimulating macrophage function^39^. Lysophopshatylserine (LPS) positively regulates inflammatory responses through activation of mast cells^40^ and was down-regulated in response to antidepressant treatment. The combined effects for PA and LPS species implicate peripheral inflammation and the immune system as highly sensitive to chronic stress and amenable to regulation with antidepressant treatment. In support of this notion, there is growing evidence that inflammation and the peripheral immune system helps drive behavioral responses to stress^41^. Furthermore, we observed increases in sphingomyelin (SM) medium chain lipids (*F*_2,58_ = 5.21; *p* < 0.05; Fisher’s LSD) and ceramide monounsaturated lipids (*F*_2,58_ = 4.48; *p* < 0.05; Fisher’s LSD) specifically in the serum of susceptible animals with decreased levels of a SM species upon imipramine treatment (**Fig. 3g**). Plasma ceramide levels are elevated in individuals with MDD^42^ and many antidepressants, including imipramine, inhibit the functions of acid sphingomyelinase^43^, which catalyzes the degradation of SM to ceramide. These data collectively highlight a possible role for peripheral lipids as markers and mediators of MDD.

Lastly, this dataset offers a valuable resource in identifying circulating peripheral markers of stress susceptibility or resilience. High levels of the serum metabolites flavin adenine dinucleotide (FAD) and uric acid, among others, are associated with susceptibility, and low levels of bilirubin are associated with resilience. Interestingly, uric acid and bilirubin have been associated with MDD in humans^44,45^, indicating the translational potential of these data. The lipids 16-HETE and 12-oxoLTB4 are products of arachidonic acid and are elevated in the serum of susceptible animals (**Supp. Table 1c**), and the proteins Klkb1, Trip11, and Mug1 are decreased (**Supp. Table 1d**). Detection of a molecular profile that spans the described classes of molecules may improve reliability, and is an important area of future efforts. Recently, circulating levels of acetylcarnitine, which is an epigenetic regulator and participates in beta oxidation, was reported to be diminished in individuals with MDD^46^. While circulating levels did not reach statistical threshold in our dataset, we did observe acetylcarnitine as significantly increased in the vHipp of susceptible animals and increased in the NAc of resilient animals, reinforcing our findings on the opposing and brain region-specific contributions to CSDS-induced abnormalities.

In sum, our datasets offer new insights into the molecular mechanisms of MDD and its treatment, as we identified a range of molecular changes induced in serum and specific brain regions after CSDS and in response to imipramine administration. Further analysis of these datasets may guide efforts to construct reliable peripheral markers of MDD and its treatment and uncover novel molecular pathways for therapeutic discovery efforts.

## Supporting information

## ACKNOWLEDGEMENTS

We thank Jill K Gregory, CMI, FAMI, Associate Director of Instructional Technology at the Icahn School of Medicine at Mount Sinai for her help in formatting the metabolic network figure. This work was supported by the National Institute of Health grants K99DA045795 to PJH, and P50MH096890 and R01MH051399 to EJN, and the Hope for Depression Research Foundation.

